# Top-Down Effects on Translucency Perception in Relation to Shape Cues

**DOI:** 10.1101/2024.11.13.623357

**Authors:** Takehiro Nagai, Hiroaki Kiyokawa, Juno Kim

## Abstract

It is well established that object shape perception significantly influences the perception of translucency. However, how object shape cues such as motion and binocular disparity affect the perception of translucency in rich environments, like virtual reality or real visual environments, remains unclear. This study aims to psychophysically measure the extent to which multiple object shape cues influence the perception of translucency. Additionally, we examined whether top-down factors, such as changes in cognitive attitude caused by the sequence of experiments, affect translucency perception. The results revealed that while motion and binocular disparity enhance translucency perception, this effect is confined to situations where shape cues are poor. Moreover, the effect became particularly pronounced when the experiments began with weak specular reflection stimuli, followed by the experiments using stimuli with specular reflection. In the case of translucent objects without specular reflection, strong shape information cannot be derived solely from shading patterns. These findings thus suggest that top-down factors related to shape modulate the influence of shape cues on translucency perception.

## 1. Introduction

When humans observe objects, they form various impressions about the material properties. This process is referred to as "material perception" in this study. Unlike simple visual attributes such as shape and color, material perception involves the complex interactions between an object’s material, shape, and lighting, which generates diverse patterns in the retinal image [1]. Therefore, as with surface color perception, it is impossible to uniquely determine a material’s properties from the retinal image alone. As a result, research on material perception has primarily focused on identifying the heuristics the visual system employs to interpret material properties from the retinal image [2,3].

The properties of glossiness, a representative material feature, have been the subject of extensive research, particularly since the 2000s. Early studies identified a relationship between two-dimensional image features and glossiness perception [4,5,6], suggesting that the heuristics employed by the visual system might be relatively simple. However, later studies revealed that glossiness perception results from the interaction of surface image patterns with various factors specific to specular reflection, such as motion flow, binocular disparity, and other shape information [7,8,9]. These findings indicate that glossiness perception arises from a complex combination of multiple factors. It is likely that such multifactorial influences apply to other material features as well.

Research on the perception of translucency has been gradually increasing. While glossiness primarily reflects the surface’s reflective properties, translucency pertains to material perception associated with the light-transmitting characteristics of the material. Similarly, transparency, another perception related to light transmission, has been extensively studied, particularly in relation to object-background interactions, as exemplified by Metelli’s law [10]. In contrast, translucency perception emerges from the retinal image created by complex interactions between lighting, the object’s optical properties, and its shape. Consequently, much like glossiness, several studies have shown that complex spatial patterns of luminance and chromaticity serve as cues for translucency [11,12,13,14]. This can be attributed to the unique patterns caused by subsurface light scattering in translucent objects, which differ from those in opaque objects.

A key feature of translucency is its strong dependence on object shape. This is because translucency involves the perception of light transmission and scattering; whose appearance of object surfaces is directly influenced by the object’s three-dimensional shape. For example, when an object has both thin and thick parts, the direction of the light source has a greater impact on the sense of translucency [15]. Moreover, it has been shown that the combination of shape perception and luminance patterns is a powerful cue for evoking the perception of translucency [16]. While motion and binocular disparity are commonly used as cues for shape perception, the presence of specular highlights also provides strong shape information [17,18]. This effect is thought to be particularly strong in translucent objects, as subsurface scattering reduces the contrast of luminance patterns on object surfaces [19], making it difficult to derive shape information from shading alone [20]. Indeed, when specular highlights are present, shape perception of translucent objects improves, and perceptual translucency is enhanced as well [21].

How does visual information related to shape contribute to the perception of translucency? For instance, in studies demonstrating the influence of shape on translucency, it was shown that even when object stimuli possess identical luminance patterns, significantly altering the perceived shape through binocular disparity can affect translucency perception [16]. This strongly suggests that shape perception plays a crucial role in translucency. However, in more typical situations where various cues, such as object shape contours, are available, the ambiguity of shape perception in translucent objects is relatively low. A straightforward hypothesis is that the perceived shape, resulting from the integration of all available cues, combines with luminance patterns to produce the perception of translucency. In such cases, even a single effective cue for shape perception may suffice to significantly impact translucency perception. Furthermore, if shape recognition is indeed critical, top-down factors related to shape recognition could potentially alter the perception of translucency, even when the visual stimuli remain identical. These insights are likely important for applications such as replicating translucency perception accurately in Virtual Reality (VR) environments.

This study aims to elucidate how multiple object shape cues interact and contribute to the perception of translucency. We focus on factors relevant to VR—specifically, the strength of specular highlights, object motion, and binocular disparity—and investigate their complementary effects on translucency by systematically controlling their intensity or presence in psychophysical experiments. Additionally, to explore whether shape recognition exerts a top-down influence on translucency perception, observers were divided into two groups with different stimulus presentation sequences, and the relationship between shape cues and translucency was compared between the groups. In one group, the experiment started under conditions with strong specular highlights, where shape cues were abundant. In the other group, the experiment began under conditions with weak specular highlights, where shape information remained insufficient despite the presence of object motion and binocular disparity. Since both groups observed the same experimental stimuli, any differences in their translucency evaluations should be attributed to differences in shape recognition arising from the order of presentation.

## 2. Methods

### 2.1. Observers

Ten undergraduate and graduate students from the Institute of Science Tokyo (two of whom were women aged 22-24) participated in the experiment. All observers had normal or corrected-to-normal vision and were unaware of the experiment’s purpose. As detailed later, observers were randomly assigned to two groups of five, with each group following a different experimental order. The experiment was conducted in accordance with the Declaration of Helsinki and approved by the Ethical Review Committee of Institute of Science Tokyo. The observers were recruited between May 7, 2024, and June 11, 2024, and all experiments were completed within this period.

### 2.2. Apparatus

The experiment was conducted in a simple darkroom equipped with blackout curtains. An LCD display (CG279X, EIZO, Japan) was used, featuring a spatial resolution of 2560 × 1440 pixels and a frame rate of 60 Hz. The display’s color gamut was set to sRGB with a white point at D65, a maximum brightness of 120 cd/m², and a gamma of 2.2, calibrated using the display’s self-calibration function. These settings were verified using a spectroradiometer (Specbos 1211-2, JETI, Germany). A custom-made mirror stereoscope was positioned in front of the display, allowing stimuli from the left and right halves of the screen to be directed to the left and right eyes, respectively. The optical viewing distance to the display through the stereoscope was 50.6 cm. Observers’ heads were stabilized with a chin rest for consistent fixation.

The entire experimental procedure was controlled by a computer (GV301Q, equipped with an AMD Ryzen™ 7 5800HS processor and an NVIDIA® GeForce® GTX 1650 GPU, ASUS, Taiwan). The experiment was managed using a custom program developed with the Coder component of PsychoPy [22]. Observers provided their responses using a trackball.

### 2.3. Stimuli

Fig 1 presents an example of the stimulus. The stimuli consisted of either still images or videos on based computer graphics (CG). Below the stimulus, a black bar (R, G, B = 0, 0, 0) and a small circular cursor were displayed, allowing observers to respond using the Visual Analog Scale (VAS). The background was a uniform gray with a luminance of 32.7 cd/m² and a D65 chromaticity.

**Fig 1:**
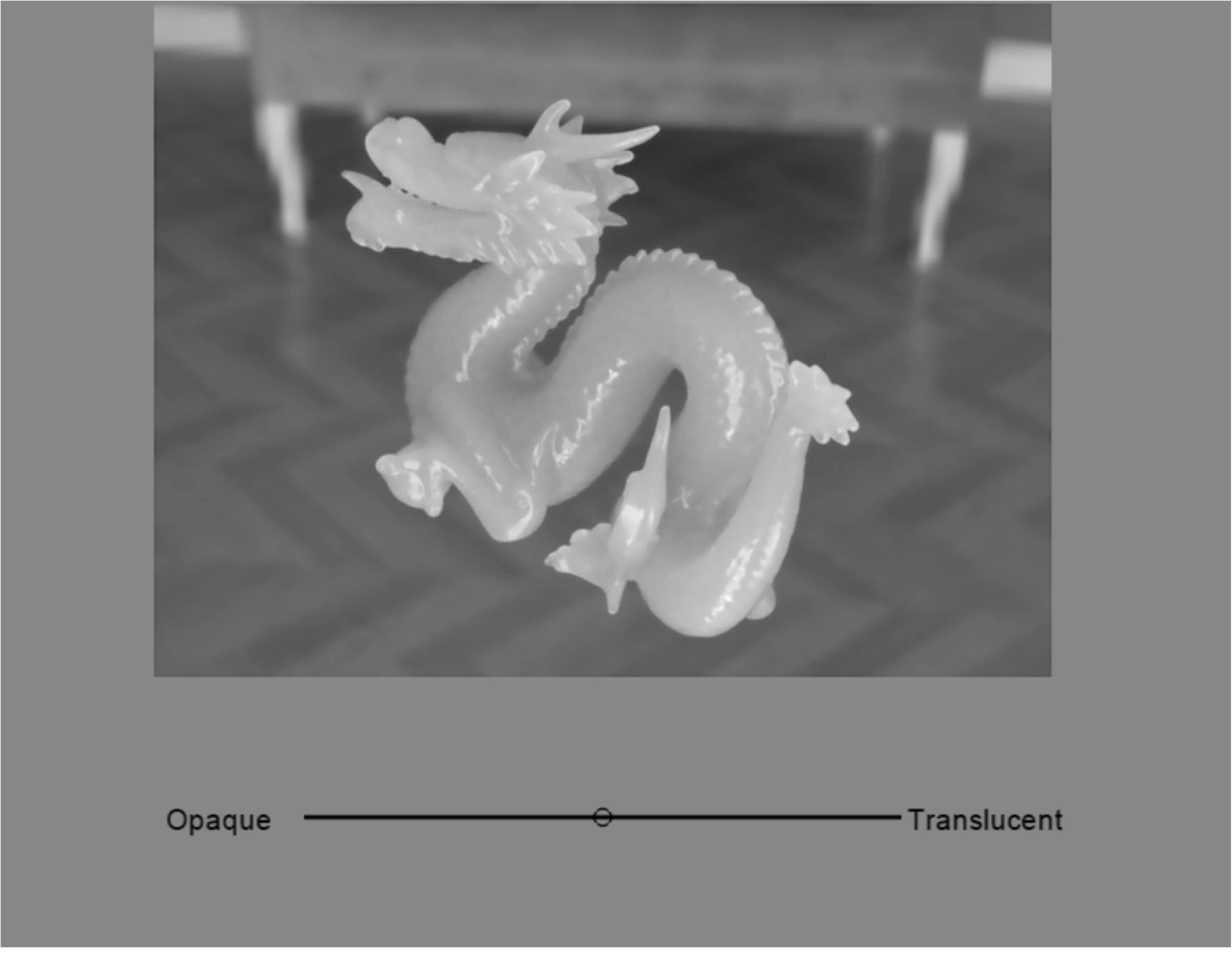
Example of a stimulus. This figure shows the stimulus presented to the left eye only. The Visual Analog Scale (VAS) scale bar and cursor at the bottom of the display were presented in both the left and right eye stimuli, with zero binocular disparity.

#### 2.3.1. Computer Graphics Rendering

The stimuli for each object were created from 800 × 600 pixel CG images rendered using Mitsuba 3 [23]. Since this study does not focus on the effect of color on translucency perception (e.g., [24]), the CG images were rendered in RGB rather than spectrum-based. An environment emitter was used for lighting, and the environment map "Brown Photostudio 06," obtained from Poly Haven (https://polyhaven.com/a/brown_photostudio_06), was employed. To create video stimuli depicting object motion, 91 images were rendered by rotating the object from -45 degrees to +45 degrees in 1-degree increments. For stereoscopic viewing, these 91 images were rendered from two separate positions, assuming an interocular distance of 6.2 cm (unit distance 62), with the camera aimed at the object placed at the spatial origin. The distance from the camera to the object’s center was set at a unit distance of 506. Specifically, the cameras were positioned at (X, Y, Z) = (±31, 385, 330) (all in unit distance).

The Bidirectional Scattering Distribution Function (BSDF) for all objects was set to *roughdielectric*, a built-in BSDF model in Mitsuba. Participating media were configured inside the objects to represent light transmission and scattering. The surface roughness parameter *α* was set to 0.05 and 0.4, corresponding to strong and weak specular highlights, respectively. For convenience, we will refer to the condition with strong highlights as the "Specular" condition and the condition with weak highlights as the "Diffuse" condition. Fig 2(a) shows examples of images rendered with these two different *α* values. The parameters for the participating media were set using a combination of the single scattering albedo (*α*) and the extinction coefficient (*σ*_*t*_). Specifically, the RGB values for *α* were kept constant, and the combinations of (*α*, *σ*_*t*_) were (0.966, 128), (0.985, 256), (0.995, 512), and (1.000, 1024). These values were chosen arbitrarily by the authors based on observations of the stimuli to induce variations in translucency. Fig 2(b) presents examples of images generated with these four combinations of *α* and *σ*_*t*_.

**Fig 2.**
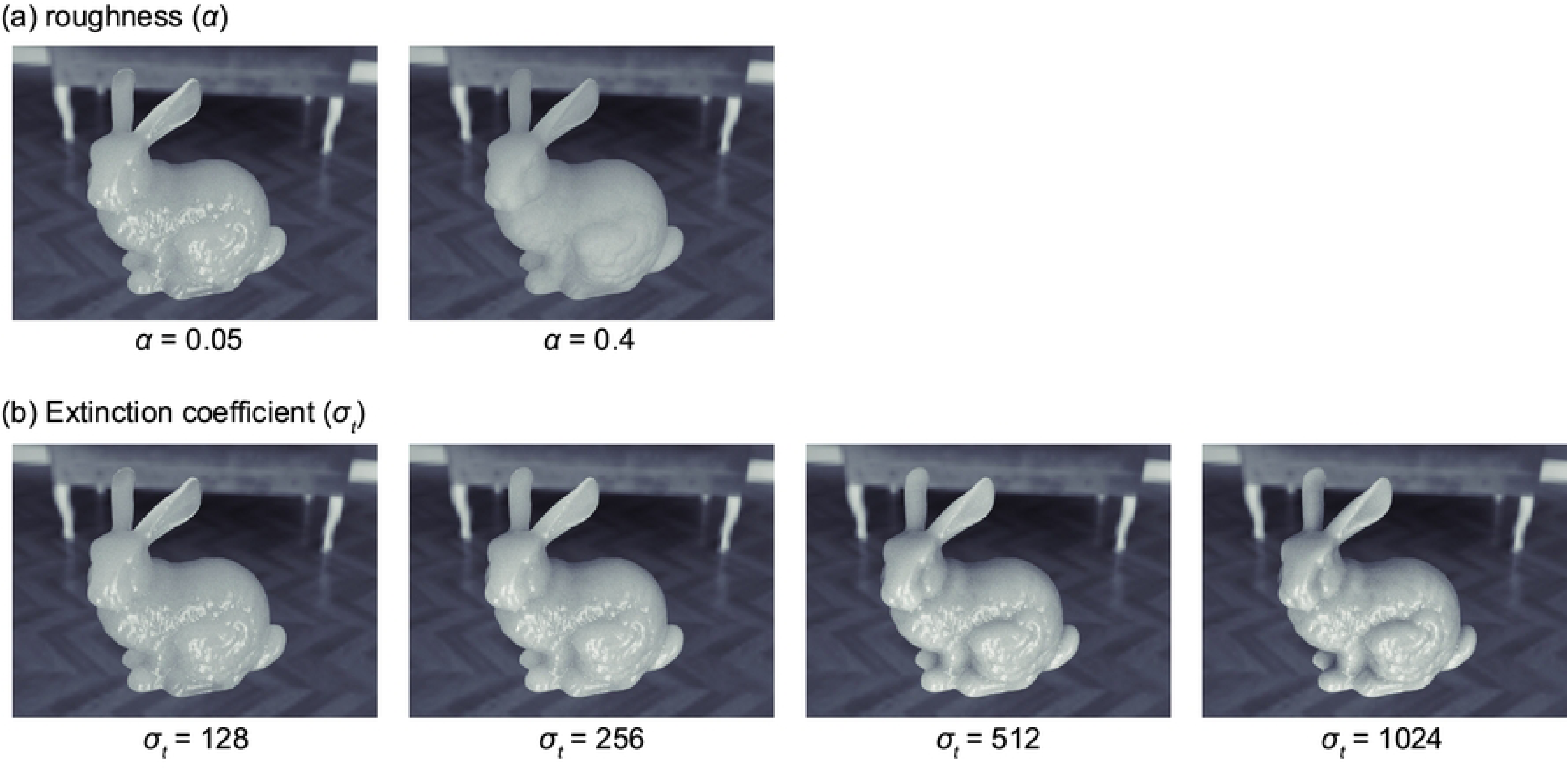
Variations in stimuli based on optical properties. (a) Changes in stimuli due to surface roughness (alpha). The left image shows *α* = 0.05, while the right image shows *α* = 0.4. (b) Changes in stimuli based on the combination of single scattering albedo and the extinction coefficient. From left to right, the extinction coefficient (*σ*_*t*_) increases.

There were four types of object shapes. Two of them, the Stanford Bunny and Dragon, were obtained from the Stanford 3D Scanning Repository (https://graphics.stanford.edu/data/3Dscanrep/). The other two shapes were blob forms created in Blender 4.1 by applying a cloud texture modifier to a UV sphere. The shape with larger surface bumps is referred to as Blob_L, while the one with smaller bumps is called Blob_S. The size of each object was adjusted to approximately 25 units in depth, width, and height. Fig 3 shows examples of CG images generated from these four shapes. Consequently, the images consisted of combinations of two surface roughness levels, four types of participating media, and four shapes.

**Fig 3.**
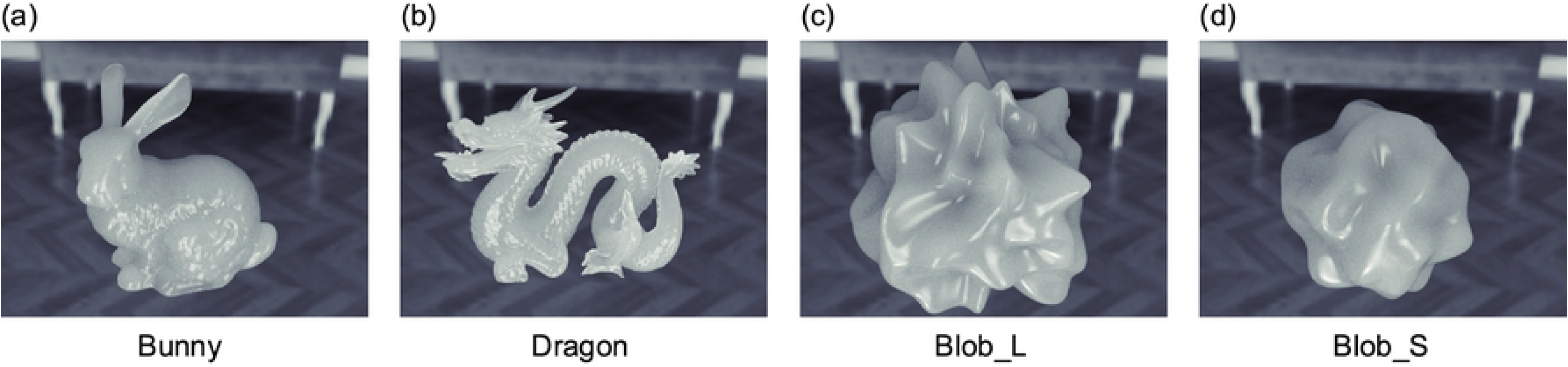
Examples of CG images generated from the four object shapes. (a) Bunny, (b) Dragon, (c) Blob_L, and (d) Blob_S.

No tone mapping was applied to the generated images. The CG images were saved as PNG files in sRGB format with a gamma of 2.2 and a D65 white point. In some pixels, the RGB values exceeded the maximum (255 in 8-bit format), and in such cases, the RGB intensity was clipped to 255. These clipped pixels were primarily located in the specular highlight areas.

#### 2.3.2. Creation of stimulus videos

Next, experimental stimuli were created using CG images. As a preparatory step, since the environment map contained colors, the rendered CG images included chromaticity information. To remove color cues, the chromaticity of all pixels was adjusted to D65 with CIE1931 (x, y) = (0.313, 0.329), while maintaining their luminance. From this point, the experimental stimuli included two variations each for contour (Full and Masked), motion (Dynamic and Static), and binocular disparity (with and without disparity, called wDisparity and woDisparity, respectively). The Full and Masked conditions were introduced to manipulate 3D shape cues, as the presence or absence of object contours significantly affects the perception of 3D shape details [20]. The procedure for creating the Full condition stimuli was as follows. First, the images were resized to 600 × 450 pixels using bicubic interpolation. Then, for each eye, a series of images from -45° to +45° was compiled into a video oscillating between these angles at 30 frames per second, saved in H264 format as an mp4 video. In the Dynamic condition, the phase of the oscillation was synchronized between both eyes and randomly determined for each trial. In the Static condition, a single random frame was extracted from the video and presented as a still image in each trial. In the woDisparity condition, the stimulus intended for the left eye was presented to both eyes, creating a zero binocular disparity condition. The visual angle of the stimulus, including the background (part of the environment map), was 15.8° × 11.9°, with the object itself measuring approximately 8.6° to 11.5° in width.

The method for creating the stimuli in the Masked condition is described here. The procedure was nearly identical to that of the Full condition, with only a minor difference in the preprocessing step. First, the stimulus size was kept at the original 800 × 600 resolution, and the chromaticity of all pixels was converted to D65. Then, all pixels outside a circle with a diameter of 240 pixels (visual angle of 6.4°), centered at (x, y) = (400, 400), were replaced with a gray background. This effectively clipped most of the object image, leaving only the central region visible and making it difficult to use contour information for shape recognition. For most objects, the circular clipped area contained only the object itself; however, in the case of the Dragon, parts of the background from the environment map were occasionally visible, depending on the object’s angle. The circular clipping was positioned slightly lower than the center to maximize the area occupied by the object within the circle.

Fig 4 presents examples of experimental stimuli for both the Full and Masked conditions. Additionally, Movie S1 and Movie S2 in Supplementary Materials show the videos for the Full and Masked conditions, respectively. Supplementary Materials also include Figs S1 and S2, which display the left-eye stimuli for all conditions.

**Fig 4.**
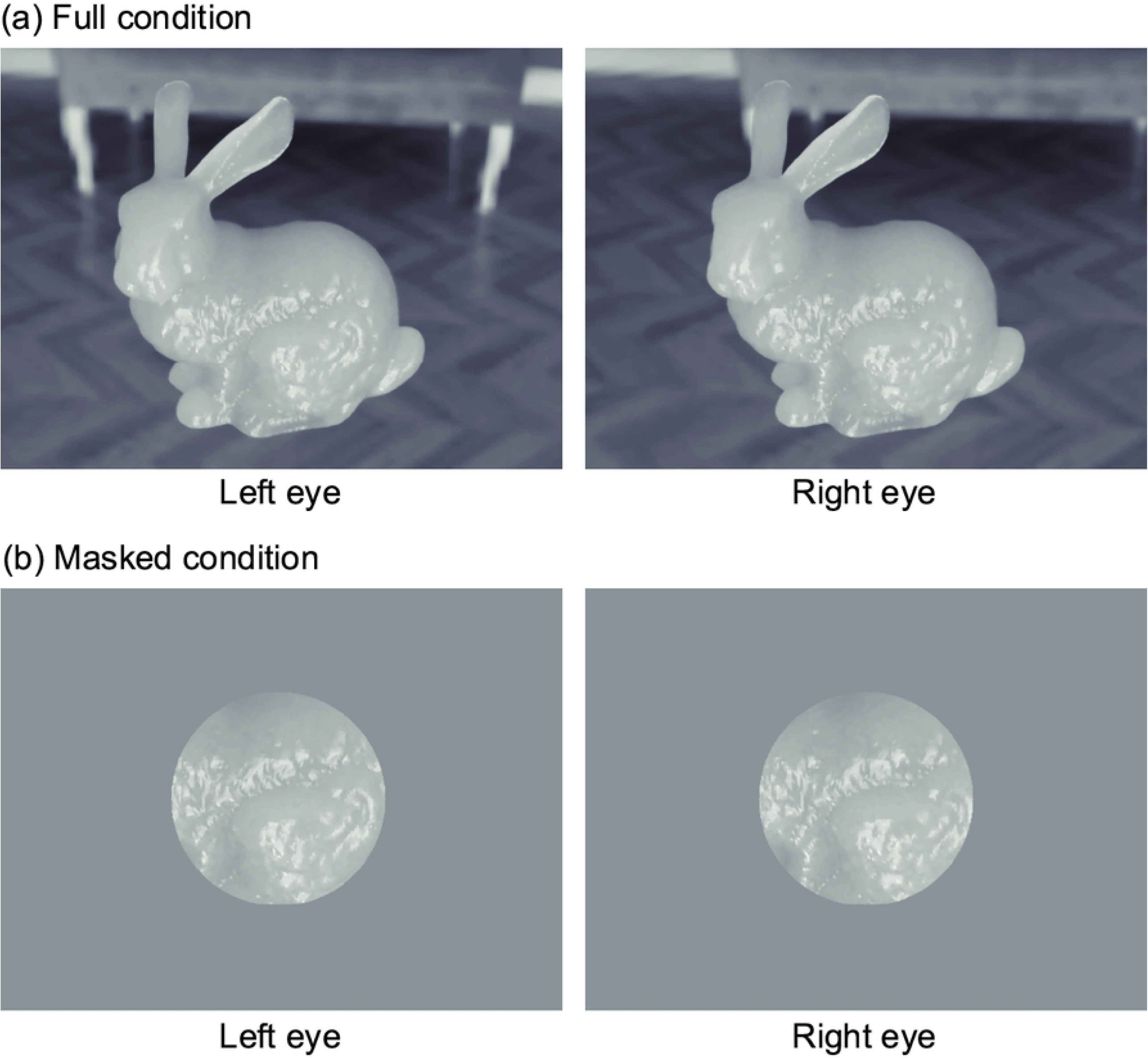
Examples of experimental stimuli. (a) Full and (b) Masked conditions. The left and right images correspond to the left and right eye stimuli, respectively. The example shown in this figure is of the Bunny under the conditions of *α* = 0.05 and *σ*_*t*_ = 128.

### 2.4. Procedure

The experiment involved rating perceived translucency using a Visual Analog Scale (VAS). In each trial, a video was presented in the Dynamic condition and a still image in the Static condition, along with a VAS response bar. Observers used a trackball to move the response cursor left or right to adjust the scale according to their perceived level of translucency, and clicked to submit their response once satisfied. The next stimulus appeared one second later. The stimuli remained visible until the observer completed their response, with no time limit imposed.

The four combinations of Specular/Diffuse and Full/Masked conditions were measured in separate sessions with two sessions per condition, totaling eight experimental sessions. Before beginning the trials in each session, observers were allowed to examine the stimulus set as much as they wanted, enabling them to establish criteria for judging translucency. Observers were instructed to use the full range of the VAS scale (from maximum to minimum) within each session and to maintain distinct judgment criteria for each session, without applying the same standards across sessions. Once observers had finalized their criteria, they informed the experimenter, and the trials began. Each session included 64 different stimuli based on four values of *σ*_*t*_, four object shapes, two motion conditions (Dynamic vs. Static), and two binocular disparity conditions (wDisparity and woDisparity). Each stimulus was presented twice, resulting in 128 trials per session. The order of 128 trials was randomized in each session.

Additionally, observers were divided into two groups of five, each following a different session order. The first group began with the Specular condition. Specifically, they completed 2 sessions of Masked-Specular, followed by 2 sessions of Full-Specular, 2 sessions of Masked-Diffuse, and 2 sessions of Full-Diffuse. Starting with the Specular condition allowed this group to obtain object shape information early, as the Specular condition provides clearer shape cues. The second group followed the reverse order of Specular and Diffuse conditions; they completed 2 sessions of Masked-Diffuse, 2 sessions of Full-Diffuse, 2 sessions of Masked-Specular, and 2 sessions of Full-Specular. In this case, particularly during the Diffuse condition, even with motion or binocular disparity, the shape cues were insufficient, leading to a reduction in perceived surface details compared to reality [20]. The first group is referred to as the "specular-first group" and the second as the "diffuse-first group."

### 2.5. Analysis

For each observer’s results, the four session types (Full-Specular, Masked-Specular, Full-Diffuse, and Masked-Diffuse) were analyzed separately due to differences in judgment criteria across sessions. First, for each session type, the data for each observer were standardized to have a mean of 0 and a variance of 1. Afterward, the group average across observers was calculated. This result is referred to as the "normalized rating." It is important to note that the normalized rating reflects only the relative translucency perception within each session and does not correspond to the absolute level of translucency across sessions. Therefore, comparisons of translucency can only be made between conditions, such as shape, *σ*_*t*_, motion, and binocular disparity, within each session type, but not between glossiness conditions or between Full and Masked conditions. Consequently, no statistical tests were conducted across session types.

Statistical analysis was conducted using a non-parametric bootstrap method. The bootstrap procedure consisted of: (1) random sampling with replacement of observers within each group, and (2) random sampling of responses within each observer, to account for both between-observer and within-observer variance. The number of resampling iterations was set to 10,000. The significance level was *α* = 0.05, and unless otherwise specified, two-tailed tests were applied. In cases involving multiple comparisons, the Holm method was used to adjust the significance level, along with pairwise tests between conditions.

## 3. Results

Fig 5 provides an example by showing the normalized ratings for the Full-Specular and Masked-Specular conditions in the diffuse-first group. It is important to note that, as with all normalized ratings in this study, observers applied different judgment criteria for translucency in the Full/Masked and Specular/Diffuse conditions. As a result, translucency magnitudes cannot be directly compared across these conditions, and the vertical axis scales also differ. As expected, smaller values of *σ*_*t*_ (i.e., less absorption and scattering) corresponded to higher perceived translucency. Linear regression was performed for each graph (all four lines), and a one-sided bootstrap test was conducted to assess the significance of the slope relative to zero (*p* < 0.001 for all conditions). In all conditions, the slopes were significantly negative. This suggests that observers were able to perceive the physical property represented by *σ*_*t*_ as translucency. Additionally, the Dynamic condition showed higher translucency perception than the Static condition. Testing the difference in normalized ratings between Static and Dynamic, averaged over *σ*_*t*_, revealed statistically significant differences in both the Full-Specular (*p* < 0.05) and Masked-Specular (*p* < 0.001) conditions. This finding indicates that object motion enhances the perception of translucency.

**Fig 5:**
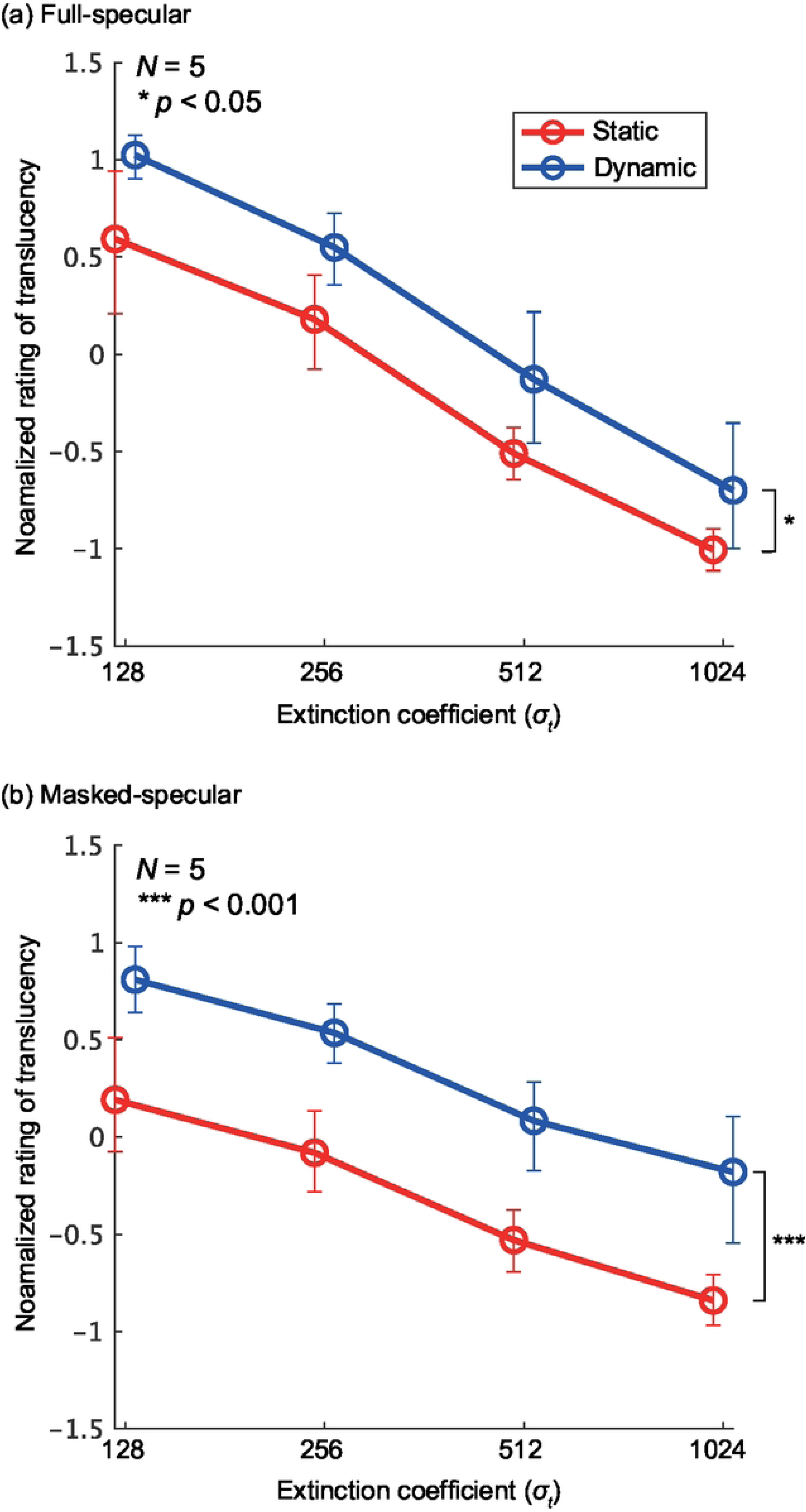
Normalized rating as a function of extinction coefficient (*σ*_*t*_) in the diffuse-first group for (a) Full-Specular condition and (b) Masked-Specular condition. The horizontal axis represents *σ*_*t*_, and the vertical axis represents the normalized rating averaged across the shape and binocular disparity conditions. The colors in the graph distinguish between the Dynamic and Static conditions. Error bars represent the 95% confidence intervals obtained via bootstrapping. Note that the vertical axis scales differ between (a) and (b), so direct comparisons of translucency between the two cannot be made.

Next, we compare the effects of motion and binocular disparity between the observer groups. Fig 6 shows the normalized ratings for the specular-first group, with the results averaged across the shapes and *σ*_*t*_. In most panels, except for the Masked-Diffuse condition, the influence of disparity and motion appears minimal. A multiple comparison test using bootstrapping was conducted to evaluate statistical differences between the combinations of motion and disparity conditions in each panel. A statistically significant difference was observed only in the Masked-Diffuse condition, specifically between the Static-woDisparity condition and both Dynamic conditions.

**Fig 6:**
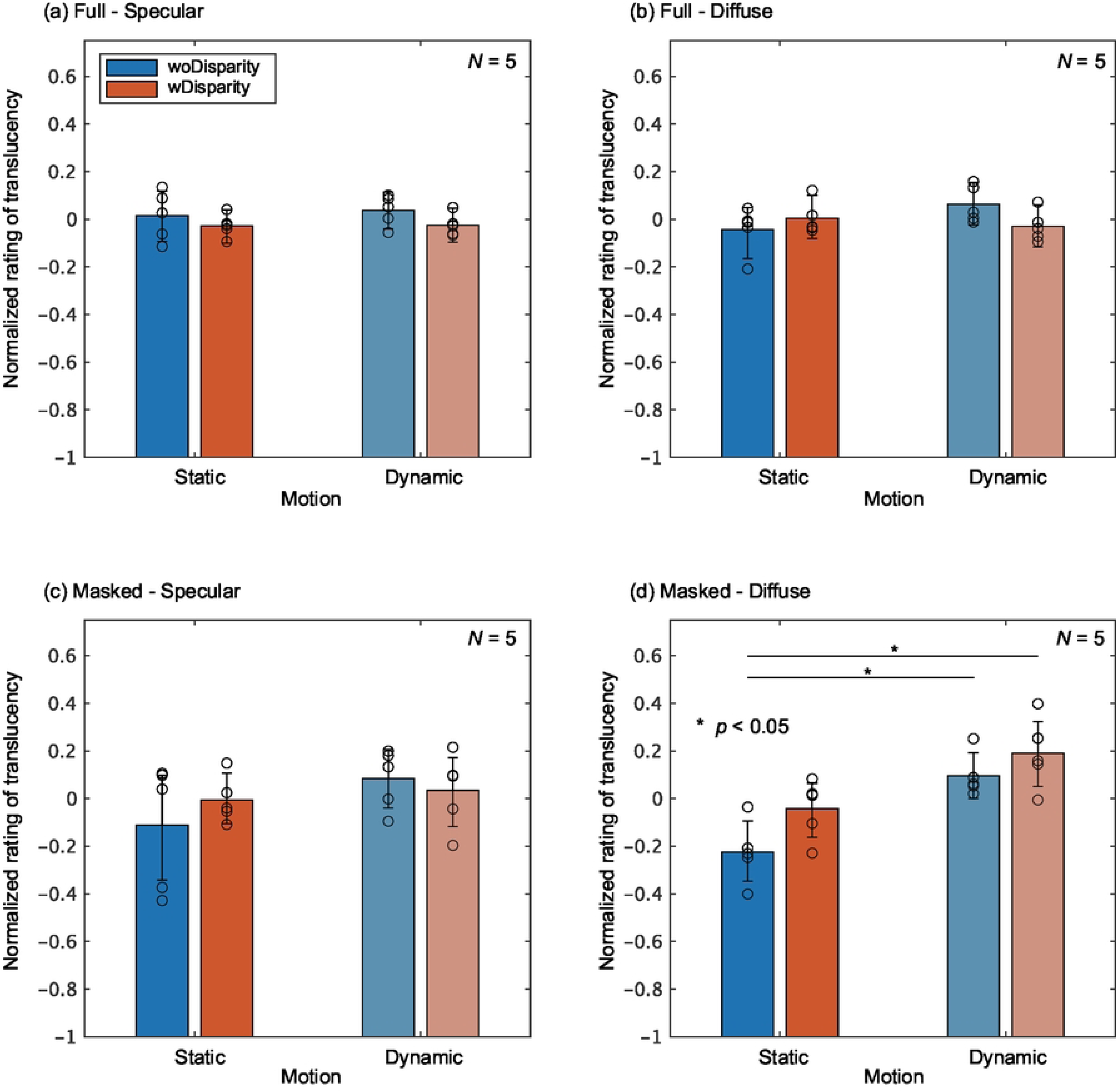
Normalized ratings averaged across shapes and *σ*_*t*_ for the specular-first group in the following conditions. (a) Full-Specular, (b) Masked-Specular, (c) Full-Diffuse, and (d) Masked-Diffuse. In each graph, the color of the bars represents the combination of motion and binocular disparity conditions. The vertical axis displays the normalized rating, with bar heights indicating the group mean and circular markers representing the results of individual observers. Error bars denote the 95% confidence intervals, which were estimated through bootstrapping. Statistically significant differences between conditions are indicated by asterisks.

Fig 7 presents the normalized ratings for the diffuse-first group. In this group, across all panels, the Dynamic condition consistently resulted in higher translucency perception than the Static condition, and wDisparity led to higher translucency than woDisparity. Similar to the specular-first group, statistical tests were conducted to evaluate the differences between motion and binocular disparity conditions. In the Full-Specular and Masked-Specular conditions, as indicated by the asterisks in Figs 7(a) and 7(c), many conditions exhibited statistically significant differences (all *p* < 0.01 or *p* < 0.001). Although no statistically significant differences were observed in the Full-Diffuse and Masked-Diffuse conditions, the Static-woDisparity condition produced the lowest normalized ratings in both cases.

**Fig 7:**
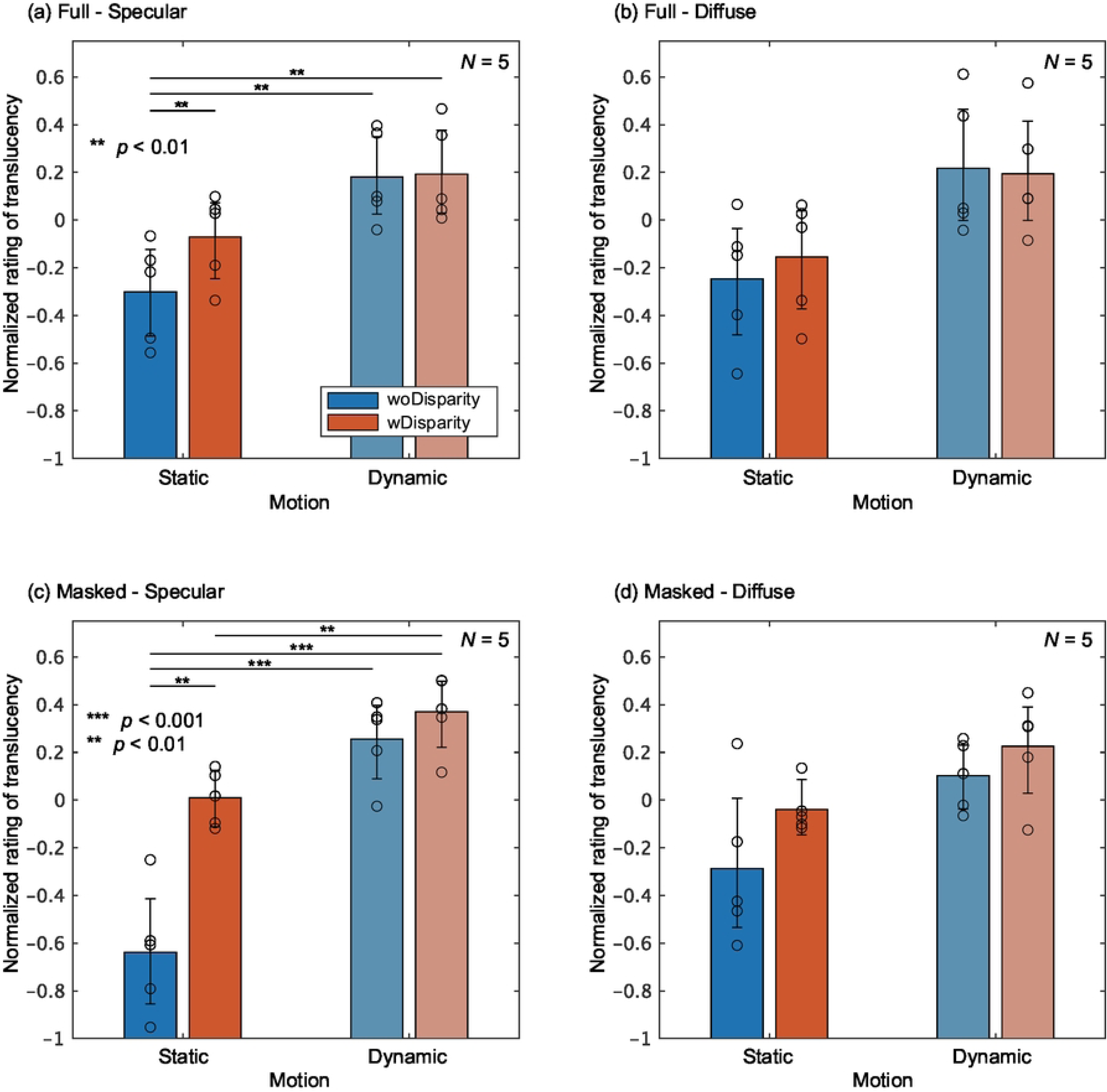
Normalized ratings averaged across shape and *σ*_*t*_ for the diffuse-first group. The format of the graphs is the same as in Fig 6.

As shown, the effects of disparity and motion appear to differ between the groups. To quantify and compare these effects, we first calculated the difference in normalized ratings between the Dynamic-wDisparity and Static-woDisparity conditions, averaged across shape and *σ*_*t*_. This difference captures the combined impact of motion and binocular disparity on translucency perception, which we refer to as the “MD (Motion & Disparity) effect.” Fig 8 shows the MD effects for both observer groups across all conditions. For the specular-first group, the MD effect was close to zero in all conditions, and no statistically significant difference from zero was found using a one-sided bootstrap test, except in the Masked-Diffuse condition. In contrast, the diffuse-first group showed positive MD effects in almost all conditions, with statistically significant positive effects in the Full-Specular and Masked-Specular conditions. When comparing the groups, the MD effect appeared larger for the diffuse-first group than for the specular-first group in all conditions. Statistically significant differences between the groups were found in the Full-Specular and Masked-Specular conditions. These results suggest that the effects of motion and binocular disparity are stronger in the diffuse-first group, particularly when the object has strong specular reflection components.

**Fig 8:**
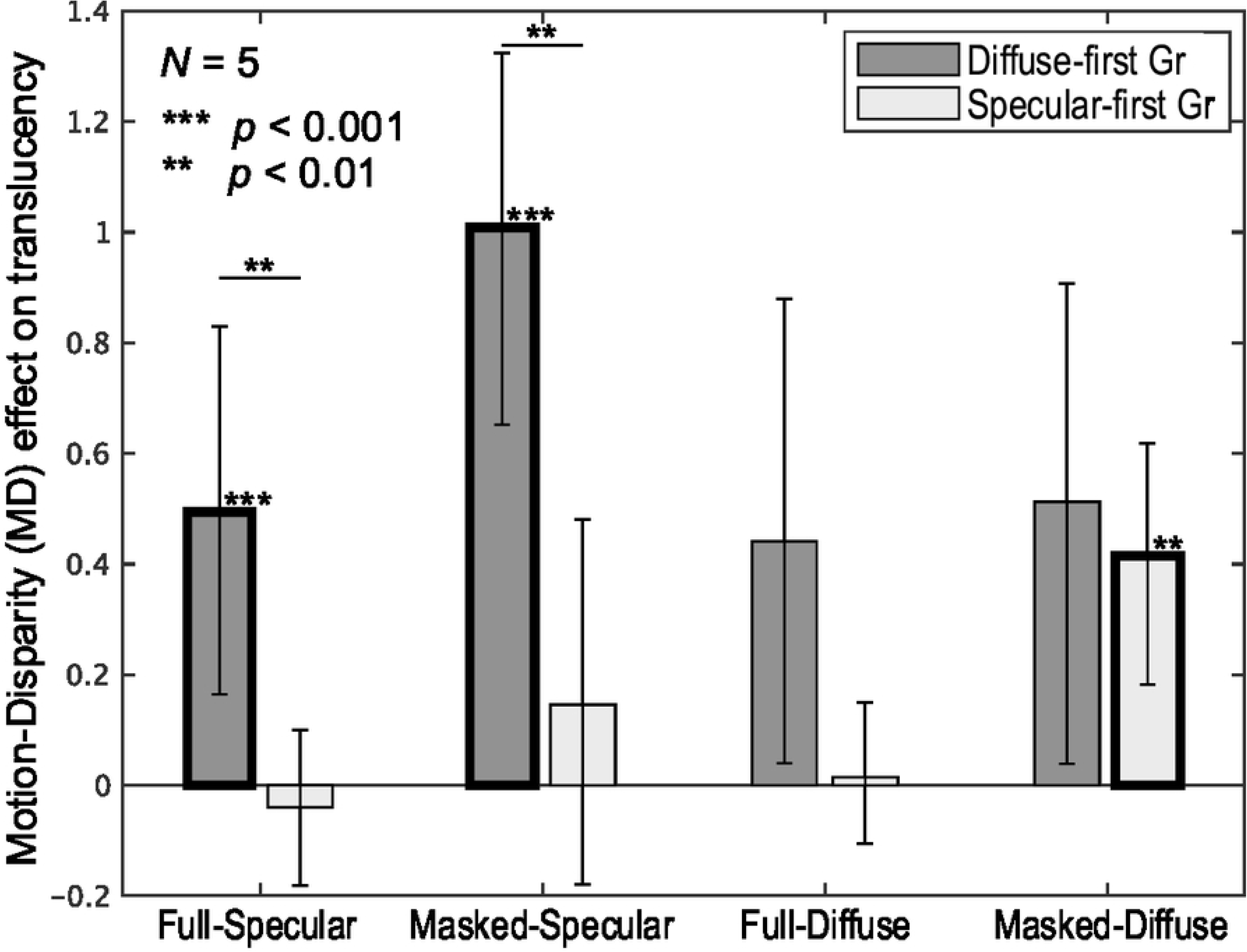
MD effect. The horizontal axis represents the combination of Full/Masked and Specular/Diffuse conditions, while the vertical axis displays the MD effect. The bar colors correspond to the observer groups. Error bars denote the 95% confidence intervals, estimated via bootstrapping. Asterisks above the bars and bold outlines indicate that the MD effects for those conditions are statistically significantly greater than zero. Asterisks spanning both the Diffuse-first and Specular-first groups indicate a significant difference in MD effect between the two groups. The results are averaged across shape and *σ*_*t*_.

Lastly, to gain insights into the mechanisms underlying the effects of motion and binocular disparity, we compared these effects across different object shapes. Fig 9 presents the normalized ratings for each object shape. As examples, the Masked-Specular condition for the diffuse-first group and the specular-first group are shown in Figs 9(a) and 9(c), and the Full-Diffuse condition results are shown in Figs 9(b) and 9(d), respectively. These two conditions were selected because they represent the largest and smallest MD effects in the diffuse-first group, as indicated in Fig 8.

**Fig 9:**
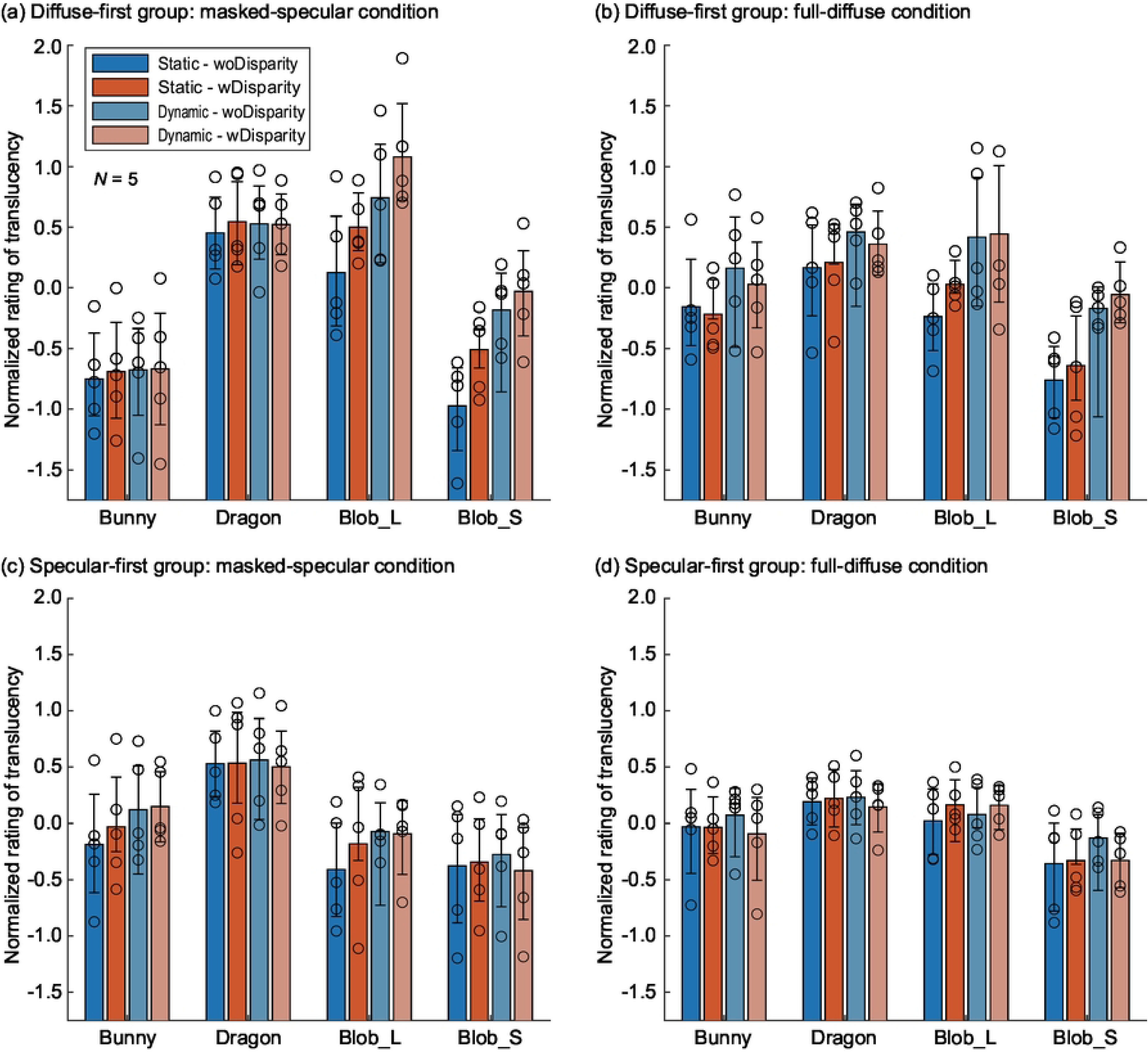
Normalized ratings for each object shape. (a) Masked-Specular condition for the diffuse-first group, (b) Full-Diffuse condition for the diffuse-first group, (c) Masked-Specular condition for the specular-first group, and (d) Full-Diffuse condition for the specular-first group. In each panel, the horizontal axis represents the object shape, and the vertical axis shows the normalized rating. The color of the bars corresponds to the motion and disparity conditions. Bar heights represent the group average, while circular markers indicate individual observer results. In all panels, the results are averaged across the Extinction coefficient. Error bars represent the 95% confidence intervals estimated via bootstrapping.

Several key points can be drawn from this figure. First, across all graphs, the normalized ratings for the Bunny and Dragon shapes show minimal differences between the motion and disparity conditions. In contrast, for the diffuse-first group, the normalized ratings for the two Blob shapes increase with the addition of each cue (motion and binocular disparity). When the MD effect (the difference in normalized ratings between Static-woDisparity and Dynamic-wDisparity) was tested for each shape, the Full-Diffuse condition, which had a small overall MD effect, showed statistically significant differences only for the two Blob shapes (*p* < 0.05 for Blob_L and *p* < 0.001 for Blob_S). These findings suggest that the influence of motion and binocular disparity on translucency perception is highly dependent on object shape.

Next, for Blob_L and Blob_S, the changes in normalized ratings due to motion and binocular disparity were nearly absent in the specular-first group but were clearly evident in the diffuse-first group. In the diffuse-first group (Figs 9(a) and 9(b)), the MD effect was statistically significantly greater than zero for both Blob_L and Blob_S (*p* < 0.05 for Blob_L and *p* < 0.001 for Blob_S under the Full-Diffuse condition, as noted earlier, and *p* < 0.001 for both Blob shapes under the Masked-Specular condition). Conversely, in the specular-first group (Figs 9(c) and 9(d)), the MD effect for the same conditions was not statistically significant for both Blob shapes and both conditions. These results suggest that the diminishing effects of motion and binocular disparity due to the experimental sequence (i.e., the difference between the observer groups) are evident even in shapes and conditions where these cues typically have a substantial impact on translucency perception.

## 4. Discussion

In this study, we examined the extent to which motion and disparity cues, which provide 3D shape information, contribute to the perception of translucency. The experimental conditions included two motion types (Static and Dynamic) and the presence or absence of binocular disparity, both of which influence shape perception. Additionally, we explored whether top-down factors, such as cognitive changes induced by the experimental sequence, affect translucency perception. To investigate this, we focused on specular reflection components, which strongly convey shape information, and assessed the impact of stimulus order—starting with stimuli that have strong specular reflection (facilitating shape recognition) versus weak specular reflection (making shape recognition more difficult)—on translucency perception, using the same set of stimuli.

As shown by the MD effect in Fig 8, in many conditions, the addition of motion and binocular disparity information significantly enhanced translucency perception. This finding is consistent with previous research showing that, even when surface luminance patterns remain unchanged, motion and binocular disparity information can alter the perceived 3D shape of an object, thereby influencing translucency perception [16]. More specifically, motion and binocular disparity likely contributed to improved shape perception, which enhanced translucency perception. In objects with subsurface scattering, surface luminance contrast and the accompanying luminance gradients are reduced, diminishing the perception of surface details based on shading [20]. Motion and binocular disparity may compensate for this reduced shape perception, leading to more accurate shape recognition. Consequently, this could result in a more precise perception of translucency. These findings emphasize the importance of incorporating motion and binocular disparity cues to support accurate translucency perception.

However, it is important to note that the effects of binocular disparity and motion are highly condition-dependent. As shown in Fig 9, these effects were mainly observed in shapes like Blob_L and Blob_S, where the original shapes were difficult to estimate. In contrast, for shapes like the Dragon or Bunny, which have abundant contour information and are easier to imagine, there was little to no effect. Additionally, as shown in Fig 8, the MD effect tended to be stronger in the Masked condition, where contour information could not be used for shape perception [20], compared to the Full condition. This trend was especially prominent in the Diffuse condition for the specular-first group and in the Specular condition for the diffuse-first group. Therefore, the effects of binocular disparity and motion are most pronounced when object shape is difficult to perceive. On the other hand, when the shape is already familiar or can be sufficiently inferred from contours in static images, these effects are likely minimal. In environments rich in visual information, such as everyday scenes or VR environments, where multiple 3D shape cues like object contours and self-occlusion are available, the contribution of motion and binocular disparity to translucency perception is likely to be limited.

Another intriguing result of this study is the substantial difference in the effects of motion and binocular disparity between the observer groups. Specifically, the diffuse-first group exhibited a much stronger enhancement in translucency perception due to motion and binocular disparity compared to the specular-first group. Since the stimuli were identical for both groups, with only the order of presentation varying, this suggests that certain top-down factors during stimulus observation influenced the impact of motion and binocular disparity. Notably, the difference in these effects between the groups was observed only for the Blob stimuli, implying that the top-down factor driven by the order of presentation is likely related to shape recognition.

However, the exact reason for the differences between the observer groups remains unclear. One possibility involves the influence of shape recognition. In the Diffuse stimuli, even with motion and binocular disparity, the perception of 3D bumps may have been diminished due to weakened luminance shading cues. In contrast, the Specular stimuli provided abundant shape information through specular reflection components [17,18], likely resulting in more prominent perceived shape details. Therefore, in the specular-first group, shape recognition may have been established early on, leaving little room for motion and binocular disparity to further enhance shape perception or translucency. However, this hypothesis is difficult to reconcile with the fact that differences between the groups were observed only with the Specular stimuli. In the diffuse-first group, once observers were exposed to the Specular stimuli, shape recognition should have been similarly established as well as in the specular-first group, leading to a rapid reduction in the effects of motion and binocular disparity, which might have minimized the differences between the groups. Alternatively, the modulation of recognition on perceived shape by attention or priming effects may be involved. In the diffuse-first group, observers primarily viewed diffuse stimuli—where shape perception was more challenging—during the first half of the experiment. In the second half, they encountered stimuli with strong specular highlights. This contrast may have drawn their attention to the specular highlights, which were both novel and rich in shape information, or to the clearer 3D shapes. In this case, the effect of motion in enhancing the influence of specular highlights [18] could be amplified. However, binocular disparity from specular highlights conveys depth information that differs from the object’s surface and can distort perceived depth [25], suggesting that the disparity from specular highlights did not directly contribute to shape perception. Instead, the disparity effect in the diffuse-first group may have been more closely related to shape perception based on shading cues. Also, in addition to the attentional effects, shape-related priming effects may have led to more distinct 3D shape perception in the specular stimuli, contrasting with the less defined shape information in the diffuse stimuli.

Our findings do not exclude the possibility of a mechanism for translucency perception based on two-dimensional image features. Although the relationship between two-dimensional image features and translucency perception has yet to reach a definitive conclusion, it has been investigated in numerous psychophysical studies [12,19,26,27,28]. Some studies have explored this relationship in the context of how two-dimensional image features influence translucency perception via three-dimensional shape information. For example, illusory translucency induced by specular highlights has been shown to correlate more strongly with luminance gradients related to shape than with shape perception itself [13]. Additionally, physical constraints between 3D shape and translucency suggest that these constraints may affect the perception of translucency from two-dimensional image features [21]. While our results indicate that 3D shape perception plays a significant role in translucency perception, we cannot rule out the possibility that translucency perception may arise directly from two-dimensional image features without reliance on 3D shape information. Further investigation is needed to clarify the specific processes involved in translucency perception from two-dimensional image features.

The findings of this study provide a strong example of the influence of higher-level recognition in material perception research. Much of the prior work on glossiness and translucency has focused on feedforward processing. For example, a previous study [29] demonstrated that material perception changes significantly based on the recognition of viewing distance, highlighting how feedback from higher-level recognition can influence material perception. Similarly, combining visual and auditory stimuli can dramatically alter material perception [30], and this multimodal interaction might also involve higher-level recognition. Additionally, in painting, expectations—such as the notion that "the Caribbean Sea appears more transparent"—have been suggested to affect translucency perception [31]. While many aspects of our results remain unclear, they can be understood within a similar feedback framework, which may be flexible enough to change simply based on stimulus order. Therefore, even when focusing on bottom-up processing in material perception research, it is crucial to carefully control for the effects of recognition to avoid unexpected results.

Finally, it is important to emphasize that this study did not provide direct evidence regarding the top-down information that influenced translucency perception through order effects. First, it is necessary to investigate whether the differences in translucency perception caused by presentation order are related to shape recognition. One approach would be to ask observers not only about translucency but also about their perceived shape (particularly surface bumps and indentations). Since perceived shapes may differ depending on the stimulus presentation order and between observers, this would allow for an exploration of the correlation between shape recognition and translucency perception. Measuring shape perception is frequently conducted in studies of material perception [17,21,20,32]. Additionally, manipulating visual attention to shape cues would be crucial for understanding their influence on translucency perception. Moreover, given that translucency perception spans multiple material categories, such as soap and candles, it may be constructed from several perceptual dimensions. If this is the case, it is possible that order effects influenced material recognition first, which then impacted translucency perception. This possibility could also be explored experimentally.

## 5. Conclusions

This study aimed to clarify the impact of multiple shape-related cues, such as motion and binocular disparity, on translucency perception. Additionally, we investigated whether differences in shape recognition and cognitive attitude, influenced by the experimental order, affect translucency perception through top-down processes. The results demonstrated that motion and binocular disparity do enhance translucency perception, but their effect is limited to situations where shape-related cues are insufficient. Furthermore, this effect was notably limited in observers who initially experienced stimuli with strong specular reflection, while it was significantly stronger in those who first encountered stimuli with weak specular reflection. Since specular reflection provides stronger shape cues than shading in translucent objects, these findings suggest that when shape recognition is incomplete due to prolonged exposure to weak specular reflection, top-down influences increase the importance of shape cues in subsequent translucency judgments.

